# A photoswitchable cannabinoid for precision treatment of refractory hippocampal discharges in a mouse epilepsy model

**DOI:** 10.64898/2026.05.04.720358

**Authors:** Sofie Bournons, Miroslav Kosar, Bilal Kicin, Roman C. Sarott, Evelien Hendrix, Rudolf Ganzoni, Patrick Pfaff, Tristano C. Martini, Matthias V. Westphal, Michael A. Schafroth, Gino De Smet, Carina De Rijck, Liam Nestor, Robrecht Raedt, Erick M. Carreira, Dimitri De Bundel, Ilse Smolders

## Abstract

Temporal lobe epilepsy (TLE) has an unmet need for precision treatments targeting the seizure focus while avoiding effects on other body parts to minimise side effects. Photopharmacology could enable precision treatment by combining systemic administration of a photoswitchable drug with implantation of an optic fibre in the epileptic focus to induce light-dependent drug conversion from an inactive to an active configuration that interacts with its target receptor to suppress seizures. The photoswitchable Δ^9^-tetrahydrocannabinol (Δ^9^-THC) derivative, *azo*-THC-3, transitions from an inactive *trans* to an active *cis* configuration upon UV irradiation. We demonstrate that local or systemic administration of *azo*-THC-3 and local UV irradiation in the hippocampus supresses difficult-to-treat seizures in the intrahippocampal kainic acid mouse model of TLE. Furthermore, our findings illustrate that the photoswitch strategy avoids hypolocomotion, a common side effect of systemic Δ^9^-THC administration. As such, we provide the first demonstration of seizure suppression with the systemic administration of a photoswitchable compound and its local photoactivation in the seizure focus.

**Graphical abstract:** 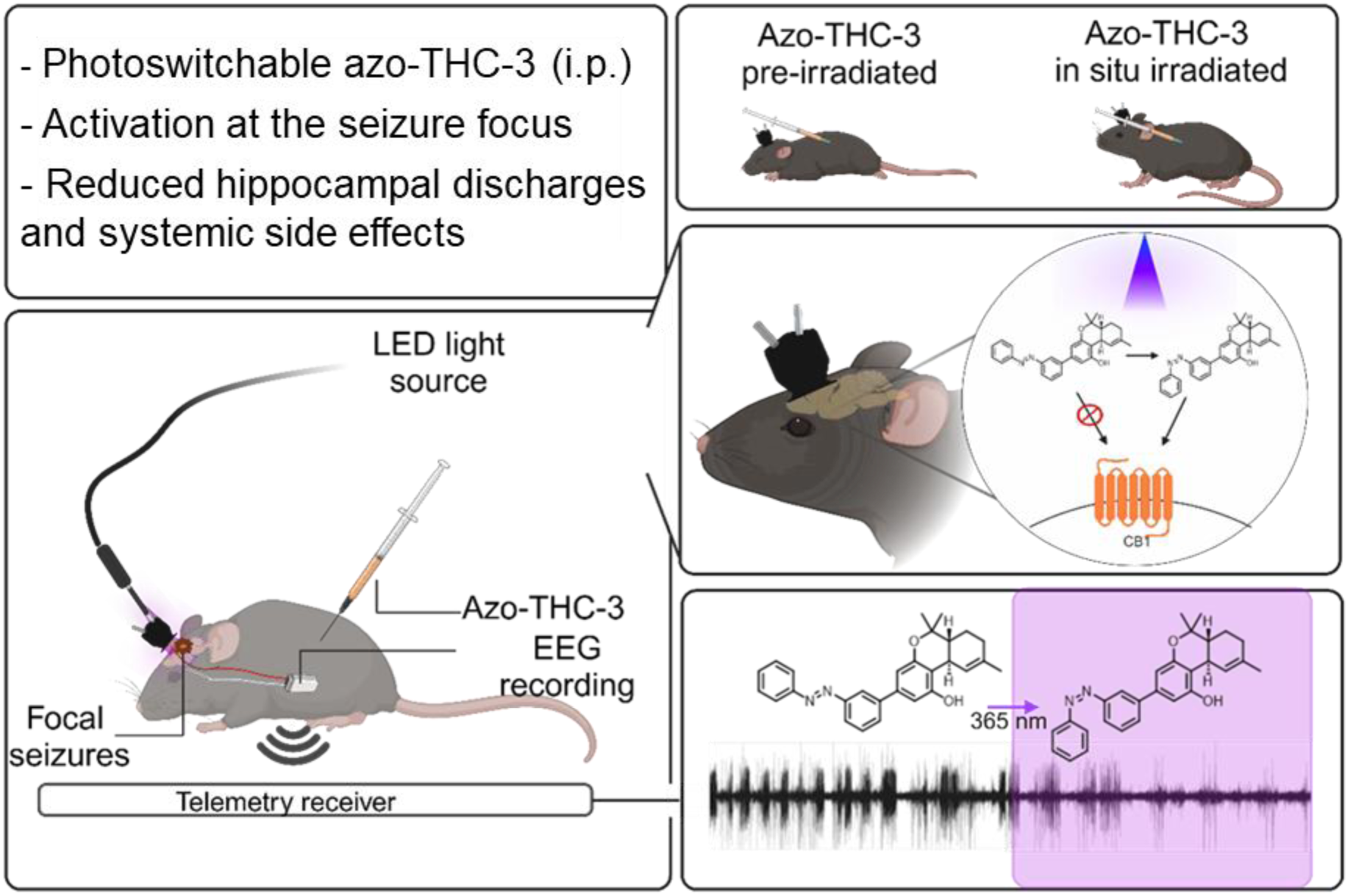

## 1. Introduction

Temporal lobe epilepsy (TLE) stands out as the leading cause of drug-resistant seizures^1,2^. It is a form of focal epilepsy with a discernible seizure onset zone in the temporal lobe of one hemisphere, often the hippocampal formation^3^. TLE seizures contribute to brain damage and are associated with psychiatric and cognitive comorbidities, reduced quality of life and elevated mortality rates, particularly in drug-resistant patients^4^. Therefore, the development of novel interventions that act specifically in the affected brain region to fully suppress seizures without major side effects remains a priority in the management of TLE. The emerging field of photopharmacology offers significant potential in this regard. Photopharmacology enables precise spatiotemporal control over therapeutic activity using focal light delivery in the brain region of interest to control the activity of systemically delivered pharmacological ligands^5–8^.

A notable photopharmacology strategy employs photoswitchable compounds that incorporate a photoresponsive moiety, typically an azobenzene. The photoresponsive motif enables light-induced transition between the distinct *trans* and *cis* configurations (see graphical abstract) and thus may mediate irradiation-dependent changes in affinity and efficacy towards the target receptor^9^. We propose that photopharmacology is a viable strategy for precision treatment of seizures in focal epilepsies such as TLE. Following systemic administration of an appropriate photoswitchable ligand in its inactive form and local light delivery through an implanted optic fibre in the brain where the seizures originate, it should be feasible to specifically switch the compound ‘on’ in the seizure focus. This should suppress spontaneous seizures while avoiding off-target as well as on-target side effects such as motor impairment due to activity of the administered compound in other parts of the body and brain. Additionally, the compound can be deactivated by applying light of a different wavelength, allowing for reversible control over its activity. To the best of our knowledge, this approach has not yet been performed in a model for TLE. Compared to other precision interventions such as optogenetics and chemogenetics, photopharmacology offers the advantage of circumventing the need for genetic engineering of brain tissue while still allowing strict spatiotemporal control over endogenous targets.

Over the last decades, the intricate role of the endocannabinoid system in regulating synaptic function has become uncovered, marking its potential for the development of novel anticonvulsive therapies^10,11^. This system consists of endocannabinoids, their metabolic machinery and the type 1 and type 2 cannabinoid receptors (CB1R and CB2R)^12^. Endocannabinoids are released into the synaptic cleft following heightened postsynaptic activity. They bind to CB1Rs, predominantly expressed on neuronal axon terminals, and attenuate further neurotransmitter release. This event may occur at both inhibitory and excitatory synapses and is termed ‘depolarisation induced suppression of inhibition’ and ‘depolarisation induced suppression of excitation’, respectively^13^. Agonists of CB1R, such as delta-9−tetrahydrocannabinol (Δ^9^-THC), the main psychoactive constituent of the cannabis plant, have been already shown to decrease overall hippocampal excitatory transmission and suppress seizures in several preclinical rodent models^14,15^. Notwithstanding this clear anticonvulsive potential of Δ^9^-THC, its ability to impair motor activity and coordination, its disruptive effects on learning, vigilance, and time perception as well as its propensity to induce delusions, hallucinations and panic reactions, has curtailed its translational value^16^. In essence, Δ^9^-THC would be a very interesting agent to use for its seizure-suppressing abilities in humans, if its action could be restricted to the seizure site of onset. We hypothesized that by application of a photoswitch strategy, it would be possible to maintain the efficacy of cannabinoids, while reducing their side effect profile. Westphal et al. (2017)^17^ designed a photoswitchable derivative of natural Δ^9^-THC by integrating an azobenzene motif into its native structure. The resulting compound, *azo*-THC-3, was demonstrated to interact with CB1R as an agonist only upon exposure to UV light at 365 nm and a concomitant switch to its *cis* configuration. By applying light, it is thus feasible to confine the robust biological action of *azo*-THC-3 to the site of light delivery.

Herein we provide the first proof-of-concept for a photopharmacological approach with a photoswitchable cannabinoid in TLE. We first validated the ability of photoactivated *azo*-THC-3 to suppress hyperexcitability in hippocampal slices. Next, we established the seizure suppressing effects of photoactivated *azo*-THC-3 in the mouse intrahippocampal kainic acid (IHKA) model for TLE characterised by typical difficult-to-treat spontaneous seizures (hence forward referred to as ‘epileptic’ mice). We studied hippocampal photoactivation following intrahippocampal infusion and following systemic administration of *azo*-THC-3 and assessed the anticonvulsive effects. Finally, we determined the effects of systemic administration and hippocampal photoactivation on motor function in the ‘epileptic’ mice.

## 2. Methods

### 2.1. Δ^9^-THC and its photoswitchable derivative *azo*-THC-3

Δ^9^-Tetrahydrocannabinol (Δ^9^-THC) was purchased at Cayman Europe (Tallinn, Estonia) and was used as a reference compound in the different experiments (end-user licence N° 001050 for narcotics/psychotropic substances granted to EFAR research group, Brussels, Belgium). The synthesis of the photoswitchable Δ^9^-THC, namely the *azo*-THC-3, was performed as previously described by Westphal et al (2017)^17^ in the Laboratory for Organic Chemistry of the ETH Zürich (Switzerland).

*Azo*-THC-3 switching properties were confirmed before experimental use in the EFAR research group at the Vrije Universiteit Brussel using high-performance liquid chromatography (HPLC). To this end, the *cis/trans* ratio of an *azo*-THC-3 solution (5.62 mM) in dimethyl sulfoxide (DMSO) was monitored and determined over a three-hour period (Figure 1). After determining the initial amount of both isomers in the starting solution, the latter was exposed to UV irradiation (365 nm, 5 mW, 50 ms pulses at 10 Hz) for 20 min using a microscope equipped with a light delivery system (coolLed pE-300 ultra, molecular devices). The *cis/trans* ratio was determined after UV exposure at 30-min intervals over a span of 90 min. Subsequently, irradiation with blue light (450 nm, 5 mW, 50 ms pulses at 10 Hz for 20 min) was conducted, and the ratio was monitored for an additional hour at 30-min intervals. Throughout the experiment, samples were kept in darkness, and the solution was maintained at 37 °C, except during irradiation periods. Verification of *cis-trans* isomerism of *azo*-THC-3 was performed using HPLC (LC-20AT HPLC pump, Shimadzu) coupled to an SPD-20A UV-vis detector (Shimadzu). Right before injection, samples were further diluted to 100 µM using DMSO. As a stationary phase, Fortis UniverSil HS C18 5 µm column was used which was kept at 60 °C in a CTO-20A column oven (also from Shimadzu). 20 µL of the samples were injected. Mobile phases A and B were 0.1% v/v formic acid (99%, ULC-MS grade, from BioSolve) in ultrapure water (acquired using a Sartorius Arium Pro purification system) and acetonitrile (HPLC grade, Acros Organics), respectively. The linear gradient profile started at 0.2% B; t = 1.5 min: 20% B; t = 4 min: 50% B; t = 6 min: 90% B; t = 10 min: 99.8%B; t = 13 min: 99.8% B; t =13.1 min; 0.2% B, t = 20 min; 0.2% B at a flow rate of 0.4 ml/min. For UV detection, a wavelength of 400 nm was used. Peaks were manually integrated using LabSolutions Lite software v5.82 from Shimadzu. *Cis*-*azo*-THC-3 eluded at 13.3 min, while *trans*-*azo*-THC-3 eluded at 14.6 min.

**Figure 1.**
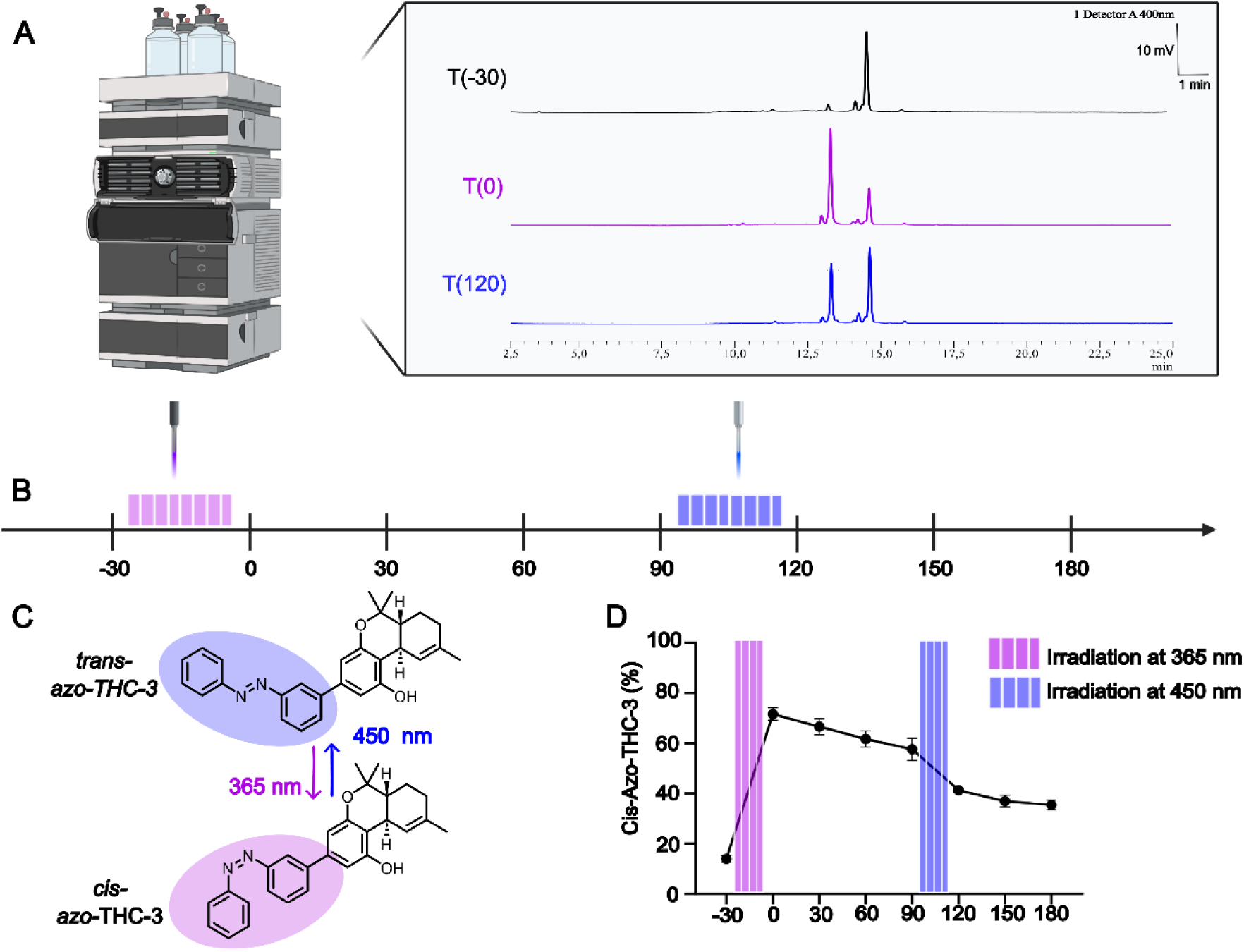
UV Irradiation of 365 nm prompts the formation of *cis*-*azo*-THC-3 but blue light irradiation cannot immediately restore the amount of *azo*-THC-3 back to initial inactive *trans*-configuration. (A) Representative chromatograms at time points T(−30 min), T(0 min) and T(120 min) after irradiation. *Trans*-*azo*-THC-3 eludes at 14.6 min whereas, *cis*-*azo*-THC-3 eludes at 13.3 min. (B) Timeline of the experiment. The injection peak within the first 2.5 minutes is not displayed. Before, and every 30 minutes after irradiation with light of depicted wavelength, the ratio of *cis/trans-azo*-THC-3 was determined. (C) Visual representation of the photoswitching properties of *azo*-THC-3. (D) The amount of *cis*-*azo*-THC-3 (% of total amount of both isomers) at the different time points following irradiation at 365 and 450 nm are presented. n = 3. Data presented as mean ± SEM. Pulsed irradiation of 50 ms at 10 Hz for a total of 20 minutes at with an intensity of 5 mW (365 nm) or mW (450 nm).

### 2.2. Mice

Experiments were conducted on male inbred C57BL/6J mice that were 7-12 weeks (Janvier, France) old at the time of arrival at the animal facility (see further section for more details). Since the objective of this study was to establish a fundamental proof-of-concept, only male mice were included to minimize inter-variability observed in the selected epilepsy model^18^. All mice were habituated to their new environment for at least one week before starting the experiments. Mice were housed under controlled conditions: the relative humidity was kept between 30%-70%, the temperature was managed between 19 °C – 23 °C and the mice were subjected to a 12/12-hour light/dark cycle. Food pellets and water were available *ad libitum*. Mice were group housed until used for brain slice experiments or until subjected to surgery. After surgery, mice were single housed for the remainder of the experiments. All experiments were authorised by the ethical committee for animal experiments of the faculty of Medicine and Pharmacy of the Vrije Universiteit Brussel (ECD 19-213-8, 20-213-1, 21-213-2, 23-213-2) and were conducted in accordance with the national (KB 29/05/2013) and European (directive 2010/63/EU) guidelines on animal experimentation.

### 2.3. Brain slice electrophysiology recordings

Twelve-week-old mice were sedated with isoflurane and decapitated. Coronal slices (300 µm thick) were prepared using a VT 1000S vibratome (Leica) in ice-cold high-choline artificial cerebrospinal fluid (aCSF) containing the following composition (in mM): 132.5 C_5_H_14_NO, 2.5 KCl, 1.25 NaH_2_PO4, H_2_O; 7 MgCl_2_.6H_2_O; 0.5 CaCl_2_.6H_2_O; 25 NaHCO_3_ and 7 glucose at 4 °C.

Subsequently, hippocampal slices were maintained at room temperature in oxygenated aCSF containing (in mM): 126 NaCl, 3.5 KCl, 1.2 NaH_2_PO_4_.H_2_0 1.3 MgCl_2_, 6H_2_O; 2 CaCl_2_.6H_2_O; 25 NaHCO_3_ and 11 glucose. The slices were then transferred to a submersion recording chamber and continuously perfused with aCSF warmed to 31 °C at a rate of 2 ml/min. All solutions were equilibrated with 95% O_2_/ 5% CO_2_. Neurons were visualized using an upright microscope equipped with a differential interference contrast optic and filter set to visualize Cornu Amonis 1 (CA1) pyramidal cells employing a water immersion objective (x40). Excitatory postsynaptic currents (EPSCs) were recorded in whole-cell configurations in voltage-clamp mode. Patch-clamp electrodes (5-8 MΩ) were filled with a cesium-gluconate intracellular solution containing (in mM): 135 gluconic acid, 135 CsOH, 10 CsCl, 0.1 CaCl_2_, 1 ethylene glycol-bis(β-aminoethyl)-N,N,Nʹ,Nʹ-tetra-acetic acid (EGTA), 10 4-(2-hydroxyethyl)piperazine-1-ethanesulfonic acid, N-(2-hydroxyethyl) piperazine-N′-(2-ethanesulfonic acid) (HEPES), 3 adenosine 5′-triphosphate magnesium salt (MgATP), 0.3 Guanosine-5’-triphosphate disodium salt (Na_2_GTP), with the pH adjusted to 7.22 using CsOH and an osmolarity of 281 mOsm/L. Neurons were clamped at 0mV, for glutamatergic events, for recording EPSCs.

During the experiment (timeline see figure 2), Δ^9^-THC and *azo*-THC-3 stock solutions in DMSO were further dissolved in aCSF and added to the bath solution to obtain a final concentration of 10µM Δ^9^-THC (based on concentration previously reported by Laaris et al.^19^) or 12.5 µM *azo*-THC-3, respectively. For *azo*-THC-3, this 25% higher concentration was used to compensate for incomplete conversion of the *trans*- to the *cis*-isomer after irradiation. Thereafter, a stock solution of kainic acid in aCSF was added to a final concentration of 150 nM (as described previously by Marsicano et al.^10^). *Azo*-THC-3 was added to the setup under dark or pre-irradiated conditions. For UV light exposure we used a UV-lamp (Universal UV lamp CAMAG type TL-900) at a wavelength of 365 nm for a minimum duration of 20 min, as was previously reported by Westphal et al (2017)^17^.

**Figure 2.**
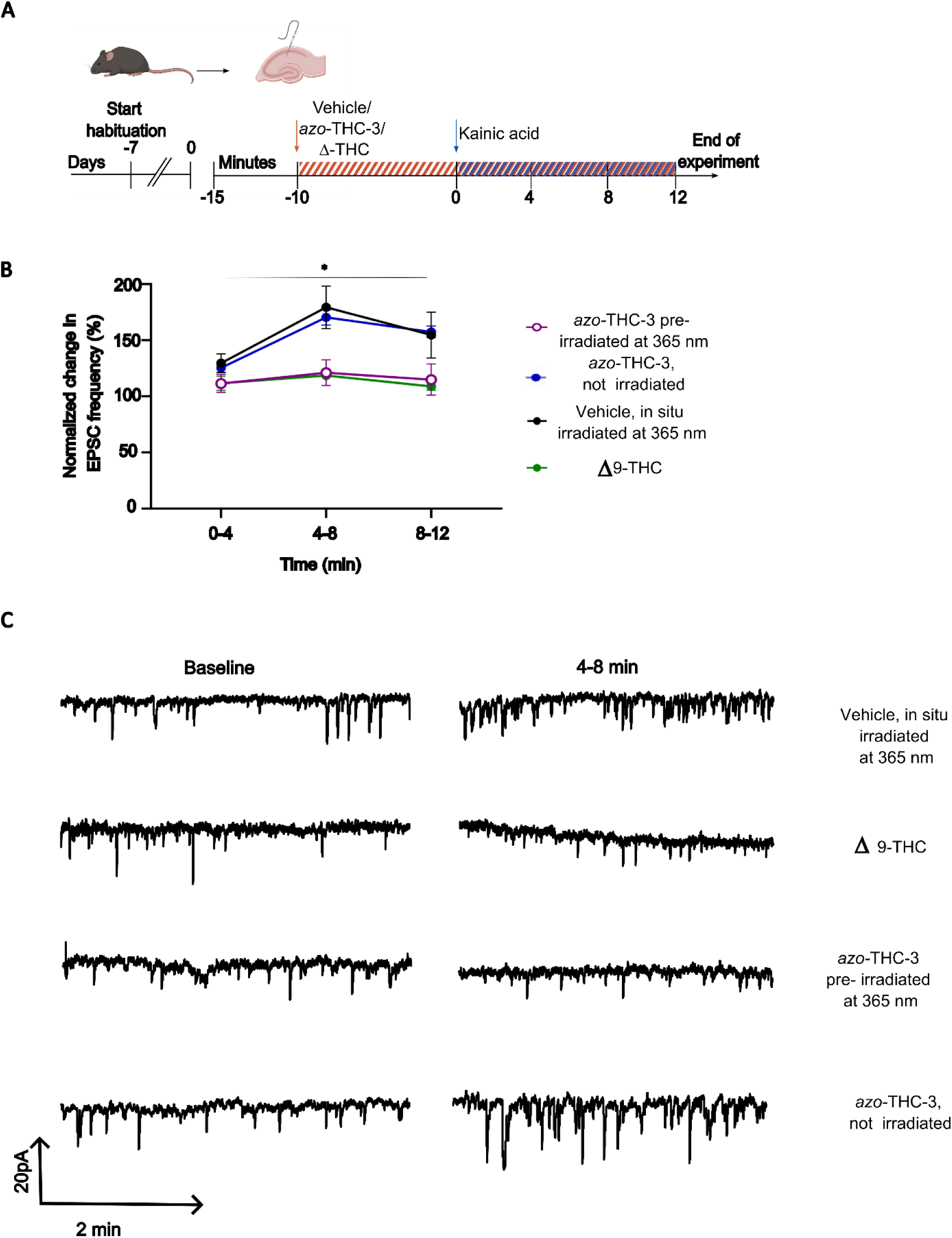
Confirmation of the light-dependent anticonvulsive effects of pre-irradiated *azo*-THC-3 in an *ex vivo* hippocampal slice electrophysiology set-up. (A) The timeline of the experiment. Patch clamp recordings were made from hippocampal pyramidal neurons from CA1 region, and four independent experimental groups were compared. Following 15-min baseline recordings, hippocampal slices were incubated with either 12.5 µM *azo*-THC-3 not irradiated, or 12.5 µM *azo*-THC-3 pre-irradiated at 365 nm, or vehicle pre-irradiated at 365 nm, or 10 µM Δ^9^−THC. After 10-min incubation, the chemoconvulsant kainic acid (150 nM) was washed into the bath solution and EPSCs were continuously recorded. (B) The time-dependent evolution of EPSC frequency is presented following the different pretreatments: *azo*-THC-3 not irradiated, n=11; *azo*-THC-3 pre-irradiated at 365 nm, n = 11; vehicle pre-irradiated at 365 nm, n = 16; and Δ^9^−THC, n = 9. Data are presented as mean ± SEM. (C) representative traces for each condition during baseline and during the second epoch (4-8 min post kainic acid administration).

### 2.4. Surgical procedures on mice

Mice that were used for the *in vivo* experiments were all first subjected to the intrahippocampal administration of kainic acid, a well validated procedure that leads to a stable amount of pharmacoresistant spontaneous paroxysmal ‘epileptic’ discharges in the hippocampal seizure focus in the chronic phase of the model (See supplementary information, Figure 1)^20,21^. To this end, mice were anaesthetised with isoflurane (Vetflurane®, 1000 mg/g, Virbac) and unilaterally injected with kainic acid, 200 ng/50 nl in 0.9% NaCl, Sigma) in the dorsal hippocampus (anterio-posterior: −2 mm; medio-lateral: 1.5 mm; dorso-ventral: −2.1 mm, relative to bregma) using a 10 µl microsyringe (65460-05, Hamilton). To prevent backflow along the injection track, the needle was left in place for 5 min. Afterwards, mice were visually monitored to ensure a status epilepticus had been induced, clearly visible by the occurrence of generalised, convulsive seizures.

After a recovery period of at least one week, a homemade probe construct was surgically implanted. This probe integrated an optofluid cannula (OmFC, with a 1.25 Zirconia Ferrule, an optical fibre core of 400 µm, a numerical aperture of 0.66, a 2 mm fibre length and a flat tip, Doric Lenses) and a stainless steel coated recording electrode (diameter: 125 µm) for measuring local intrahippocampal field potentials/electroencephalography (EEG) monitoring (E363/3/SPC, Plastics One, Bilaney Consultants) so that the recording electrode extended 1 mm below the injection cannula and the optic fibre. Employing the same spatial coordinates, the recording electrode’s tip was positioned to coincide with the site of the kainic acid injection. The probe construct was connected to a radiofrequency transmitter (ETA-F10, DSI), which was implanted intraperitoneally, and a second ground electrode was placed at the level of the cerebellum. Following implantation, mice received appropriate pain relief and prophylactic antibiotics through daily administration of Metacam^®^ (5mg/kg) and Baytril^®^ (5mg/kg) for a minimum duration of 48 h and 72 h, respectively. Four weeks after kainic acid injection, experimentation started in the epileptic mice.

### 2.5. In vivo photopharmacology experiments monitoring seizure frequency

For *in vivo* photopharmacology experiments, a Light Emitting Diode (LED)-based fibre optic system was utilised, comprising a two-channel LED driver and an LED fibre-optic rotary joint connected to an optic fibre with a diameter of 400 µm (all from Doric Lenses). *In situ* irradiation entailed delivering light pulses with an intensity of 5 mW (measured at the tip of the probe construct), each lasting 50 ms at a frequency of 10 Hz for a total of 20 min. Given the limited knowledge on the effects of UV irradiation on brain tissue, we used irradiation parameters previously reported by Zussy et al. to be well-tolerated^22^. The light delivery protocol was controlled using Doric Studio (Doric Lenses) and commenced immediately for intrahippocampal drug delivery or 5 min post-systemic injection.

Epileptic mice experience recurrent electrographic hippocampal paroxysmal discharges (HPDs), approximately 30 to 40 HPDs occur per hour, while only 1 to 2 generalised convulsive seizures are observed daily^23,24^. The substantial frequency of HPDs enables us to assess the impact of drugs by comparing them against the baseline presence. Real-time EEG recordings were acquired at a sampling rate of 500 Hz and were performed while mice were housed within their home cages, positioned on a receiver plate (RPC-1 receiver, DSI). The implanted radiofrequency transmitter (ETA-F10, DSI) transfers data to the receiver plate, which then forwards it to the data exchange matrix (MX2, Matrix 2.0, DSI), responsible for enabling communication between the telemetry implants and the Ponemah data acquisition system (DSI). The corresponding EEGs were analysed after applying a 0.008 Hz high-pass filter, a 60 Hz low-pass filter, and a 50-Hz notch filter with NeuroScore (DSI, version 3.4.0) using an automated seizure detection protocol. Automated countings were manually checked. HPDs were defined as high amplitude (> 3 times background amplitude), high frequency (> 1 Hz) sharp waves that persist for a minimum of 5 seconds. Following baseline recordings, mice were injected with either Δ^9^−THC, *azo*-THC-3 or vehicle, followed by further EEG monitoring (see Figure 3A and 4A). Compounds were administered with a minimum washout period of two days. Random drug allocation was implemented in a crossover design using Research Randomizer.

**Figure 3.**
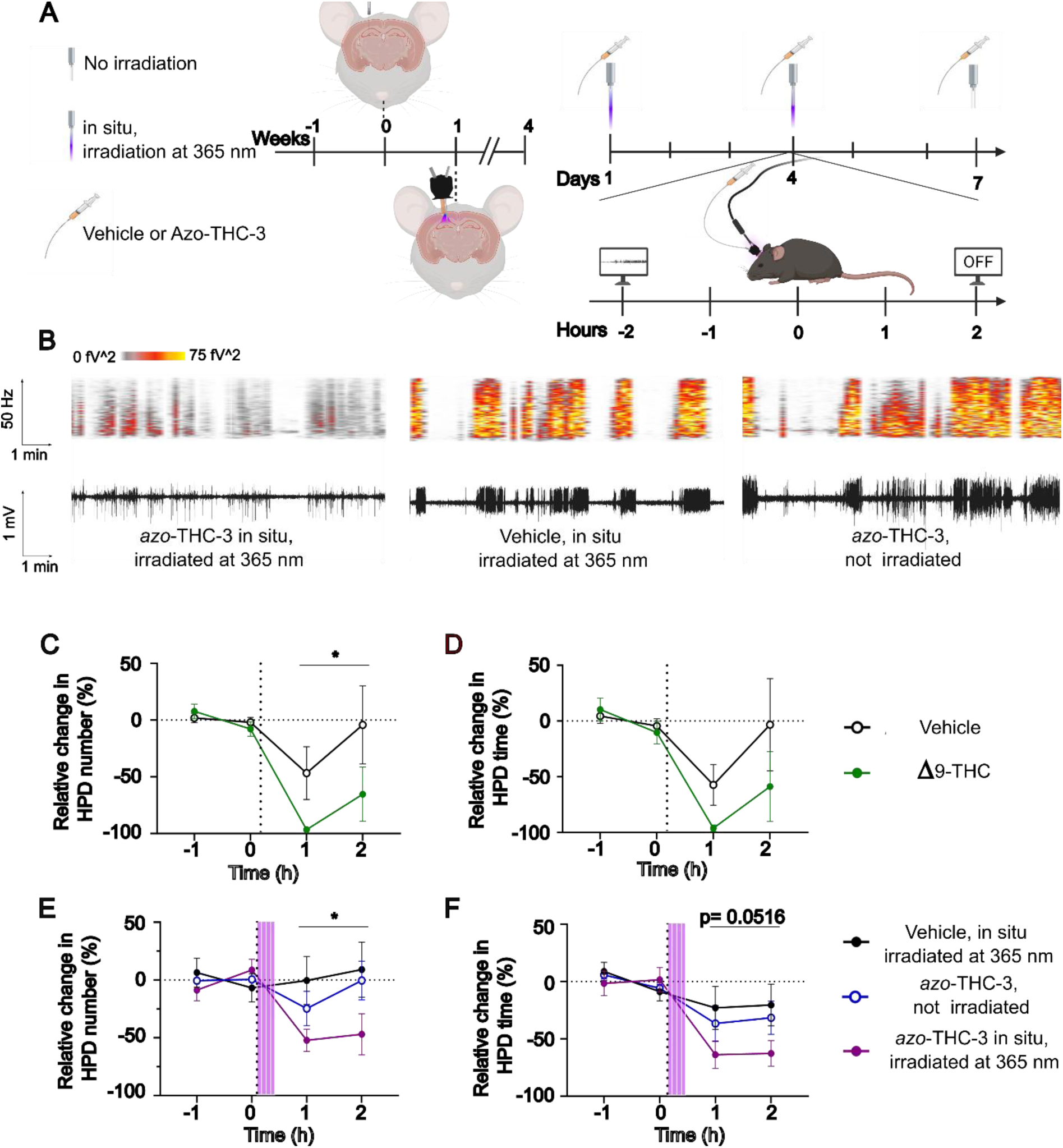
Δ^9^−THC and its photoswitchable derivative (exposed to 365 nm) reduce the HPD occurrence when locally infused in the dorsal hippocampus. (A) The timeline of the experiment. Four weeks after kainic acid injection, mice presented a stable amount of HPDs. On the day of the experiment, mice were tethered to the set-up and following a baseline period of 2 h, a drug or vehicle was infused locally. The effects were monitored for at least 2 h afterwards. (B) Ten-minute snippets of traces of the hippocampal field potentials measured *in vivo* and their accompanying periodograms 30 to 40 min post infusion for the different experimental conditions. All traces belong to the same mouse and show the difficult-to-treat HPDs typical for this chronic TLE mouse model. (C, D) The effect of vehicle and Δ^9^−THC on the number (C) and hourly cumulative timing (D). Vehicle group, n = 4; Δ^9^−THC group, n = 4. (E, F) The effect of vehicle (*in situ* irradiated), *azo*-THC-3 (not irradiated) and *azo*-THC-3 (*in situ* irradiated) on the number (E) and hourly cumulative duration (F) of HPDs. Vehicle *in situ* irradiated at 365 nm group, n = 5; *azo*-THC-3 not irradiated group, n=7; *azo*-THC-3 *in situ* irradiated at 365 nm group, n= 7. The purple stripped pattern indicates when light irradiation at 365 nm was applied for the appropriate groups and the dotted line marks the start of compound infusion. Data are presented as mean ± SEM. Overall treatment differences between experimental groups are indicated as * p < 0.05. or the exact p value is depicted.

**Figure 4.**
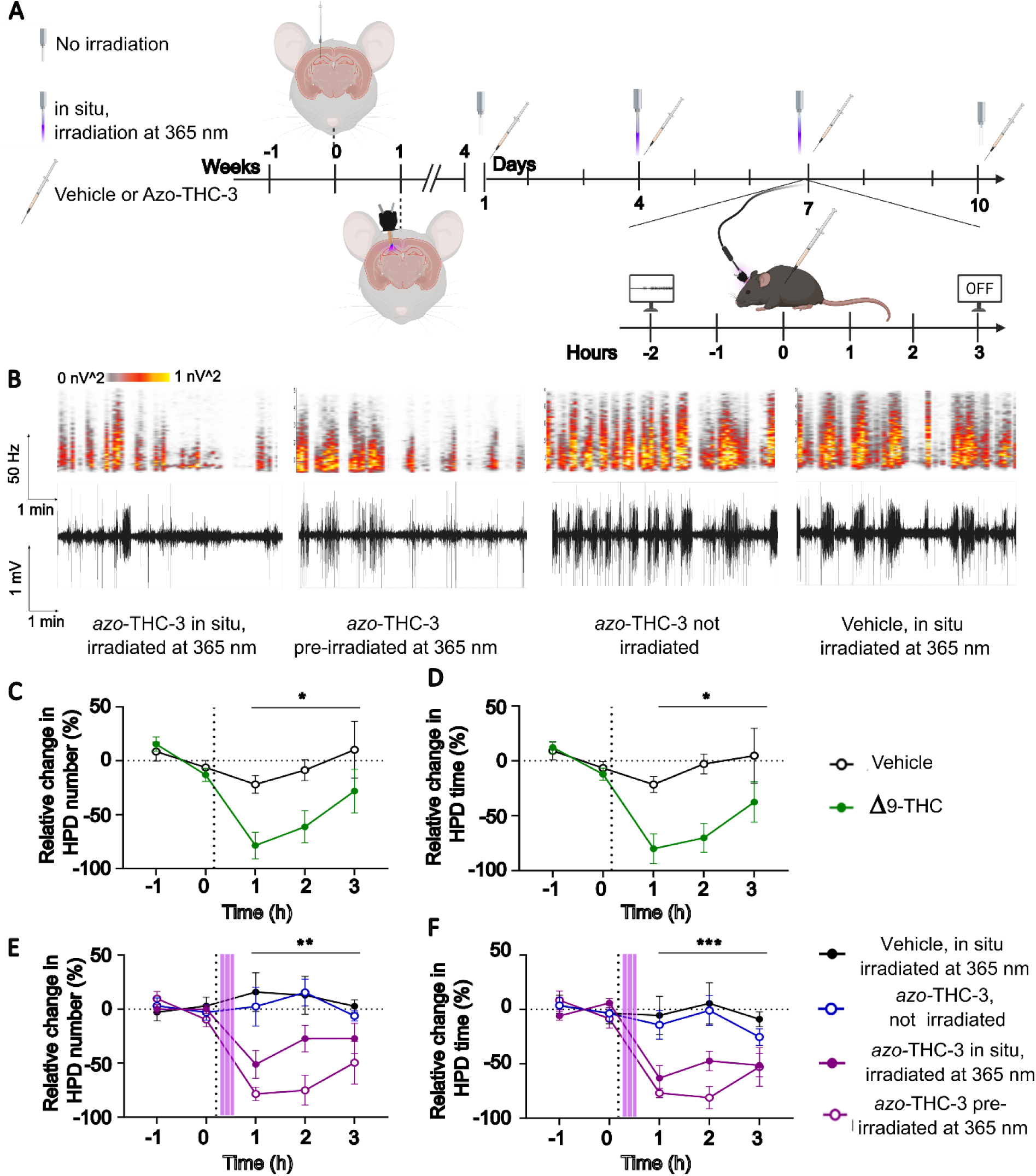
Δ^9^−THC and *azo*-THC-3 (exposed to 365 nm, pre-irradiated and *in situ*) clearly suppress the HPD burden in a chronic TLE mouse model following systemic administration of the photoswitchable compound. (A) The timeline of the experiment. Similar as in Figure 3 with as major difference intraperitoneal administration of *azo*-THC-3 compound and vehicle. (B) Ten-minute snippets of EEG-traces and their accompanying periodograms 35 to 45 min post infusion for the different experimental conditions. All traces belong to the same mouse. (C, D) The effect of vehicle and Δ^9^−THC on the number of HPDs (C) and on the hourly cumulative time of HPDs (D). Vehicle group, n = 7; Δ^9^-THC group, n = 6, (E, F) The effect of vehicle (in situ irradiated), *azo*-THC-3 (not irradiated), *azo*-THC-3 (*in situ* irradiated) and *azo*-THC-3 (pre-irradiated) on the number of HPDs (E) and hourly cumulative time of the HPDs (F). Vehicle *in situ* irradiated at 365 nm group, n = 7; *azo*-THC-3 not irradiated group, n=6; *azo*-THC-3 pre-irradiated at 365 nm group, n= 6; *azo*-THC-3 *in situ* irradiated at 365 nm, n= 8. The purple stripped pattern indicates when light irradiation at 365 nm was applied for the appropriate groups and the dotted line marks the moment drug or vehicle was injected. Data are presented as mean ± SEM. Overall treatment differences between experimental groups are indicated as * p < 0.05; ** p < 0.01, *** p < 0.001.

#### Intrahippocampal infusions

When administered intrahippocampally, Δ^9^−THC and *azo*-THC-3 were dissolved in vehicle (5% DMSO, 5% polysorbate 80, 15% polyethylene glycol 400, 75% aCSF) to a final concentration of 0.5 µg/0.5 µL or 0.96 µg/0.5 µL, respectively, on the day of the experiment. A 50 µL microsyringe (1805 RN, Hamilton) was connected to a fluid injector for multiple injections (Doric Lenses) using polyethylene tubing (Doric Lenses). Drugs were infused at a rate of 0.1 µL/min for a total of 5 min using a microinjection pump (CMA Microdialysis 400 Syringe Pump). The injector remained in place until the end of the experiment to prevent backflow along the injection track. Mice were habituated to the setup for 3 (in preliminary experiments) or 5 consecutive days before the start of the experiments to mitigate stress (see figure 3, panel C-F). *Azo*-THC-3 was infused under dark or *in situ*-irradiated conditions.

#### Intraperitoneal injections

On the day of the experiment, a stock solution of Δ^9^−THC was diluted in the vehicle (5% DMSO, 5% polysorbate 80, 90% saline) to a final concentration of 1 mg/ml and injected at a dose of 10 mg/kg. Similarly, *azo*-THC-3 in DMSO was diluted in the vehicle (5.97% DMSO, 5% polysorbate 80, 89.03% saline) and an equivalent dose to Δ^9^−THC of 23.9 mg/kg was administered. Both compounds were injected at a volume of 10 ml/kg. Prior to the experiment, mice were habituated to the setup for 3 days. *Azo*-THC-3 was injected under dark, *in situ*-irradiated or pre-irradiated conditions.

The data is presented as a relative deviation from the average amount or hourly cumulative HPD time observed during the baseline period by a scientist blinded for the different treatment groups. If, however, variation supersedes 50% within the baseline itself, the recording was excluded from analysis (for our analysis this happened in only 2 out of 69 recordings).

### 2.6. Behavioural analysis of activity and motor behaviour

We also recorded individual video files of the mouse experiments. The immobility of the mice was assessed during the initial 20-min UV irradiation/non-irradiation period (that started 5 min after compound injection) by an observer that was blinded to the different treatment groups. Immobility was defined as the lack of any movement, besides breathing.

### 2.7. Statistics

Data are presented as mean ± SEM. GraphPad Prism (version 10.0.3) is used to analyse and visualize data. All test were performed two-sided. Slice electrophysiology data is analysed using a linear mixed model (independent variables: conditions and time) followed by a Dunnett’s multiple comparison post-hoc test. The *in vivo* experiments shared a similar setup, in which a baseline recording of HPDs was challenged using different treatments. To answer the question on whether different drug conditions imparted distinct effects on HPD characteristics, a linear mixed model (independent variables: conditions and time) was performed, considering only the timepoints post drug administration. For the further behavioural analysis of immobility of the mice, ANOVA (independent variable: conditions) followed by Tukey’s multiple comparisons test was also applied. Figures were made using BioRender and Inkscape 1.3.2.

## 3. Results

### 3.1. Rapid UV-induced photoconversion of *cis*-*azo*-THC-3 to *trans*-*azo*-THC-3

We first characterized the efficacy of photoswitching between *cis*-*azo*-THC-3 to *trans*-*azo*-THC-3 at body temperature. Westphal et al (2017)^17^ reported that continuous UV (365 nm) irradiation of the *azo*-THC-3 compound for 20 min was sufficient to induce photoswitching. To avoid possible heat accumulation, we used a pulsed irradiation protocol that was previously reported to be effective and well-tolerated in mice^22^. Essentially, *azo*-THC-3 (5.62 mM) was kept at 37 °C and was irradiated at 365 nm for 20 min followed 90 min later by irradiation at 450 nm. The relative ratio of *cis*-*azo*-THC-3 to *trans*-*azo*-THC-3 was determined by HPLC at different time points (Figure 1A-C). Pulsed UV irradiation for 20 min at 365 nm resulted in an increase in *cis*-*azo*-THC-3 amount (from 13.9% ± 1.3% to 71.7% ± 2.5%) as shown in Figure 1D. Thereafter, *cis*-*azo*-THC-3 slowly relaxed back to *trans*-*azo*-THC-3. The relaxation half-life of *azo*-THC-3 at 37 °C was estimated to be 285 ± 78 min, which aligns with the value reported by Westphal et al. (2017) (t_1/2_ = 148.9 min assuming first order kinetics based on a reported τ = 216 min at room temperature)^17^. Conversely, subsequent pulsed blue light irradiation for 20 min at 450 nm did not fully revert the remaining *cis*-*azo*-THC-3 to *trans*-*azo*-THC-3 but rather induced a state consisting of 40% *cis*-*azo*-THC-3 and 60% *trans*-*azo*-THC-3. In their study, Westphal et al. observed a clear difference between UV-irradiated *azo*-THC-3 and dark-adapted *azo*-THC-3^17^. However, they did not quantify the *cis/trans* ratio. Our results show that, considering the constraints of *in vivo* irradiation through an optic fibre, a rapid UV-light induced ON switch from *trans*-*azo*-THC-3 to *cis*-*azo*-THC-3 would be achievable, whereas a rapid blue-light induced OFF switch would not be sufficiently effective.

### 3.2. UV irradiated *azo*-THC-3 prevents kainic acid-induced hyperexcitability in hippocampal slices

We next evaluated whether photoactivated *azo*-THC-3 could prevent kainic acid-induced hyperexcitability of CA1 pyramidal neurons in mouse hippocampal slices as previously described^8^. For this purpose, hippocampal slices were preincubated with either UV-irradiated 12.5 µM *azo*-THC-3 (n=11), or non-irradiated 12.5 µM *azo*-THC-3 (n=11), or our reference compound Δ^9^−THC at 10 µM concentration (n=9) or vehicle (n=16) for 10 min (Figure 2). We adapted the concentration of *azo*-THC-3 relative to the concentration of Δ^9^−THC to compensate for the difference in molar mass. Thereafter, kainic acid was added to the bath to induce hyperexcitability and the EPSC frequency was evaluated for another 12 min. Overall, the EPSC frequency increased relative to baseline levels (mixed effect analysis, time factor, F (1.732, 74.47) = 11.93, p < 0.0001). However distinct treatments affected the rise of the EPSC frequency differently (mixed effect analysis, treatment factor, F (3,43) = 3.751, p= 0.0177, interaction factor F (6,86) = 2.258, p=0.0452). As expected, Δ^9^−THC prevented this rise in EPSC frequency, with a mean difference of 60.75% (95% CI (10.30, 111.20), p = 0.0171) compared to vehicle during the second epoch (4-8 min). UV-irradiated *azo*-THC-3 was similarly effective at preventing this rise in EPSC frequency, with a mean difference of 58.23% (95% CI (2.14, 114.30), p = 0.0405) compared to vehicle during the second epoch (4-8 min). Importantly, non-irradiated *azo*-THC-3 did not affect EPSC frequencies compared to its vehicle control (0-4 min, mean difference 4.18% (95% CI (−19.61, 27.97), p=0.9496; 4-8 min, mean difference 8.99% (95% CI (−42.94, 60.91), p=0.95; 8-12 min, mean difference −2.74% (95% CI (−57.53, 52.04), p = 0.9986).

### 3.3. Intrahippocampal infusion and *in situ* photoactivation of *azo*-THC-3 reduces the incidence of the spontaneous recurrent HPDs in the seizure focus of mice

One of the main bottlenecks for the application of photopharmacology in the brain is the capacity of photoswitchable compounds to pass the blood-brain barrier and to reach the site of their desired action. Moreover, if the molecules reach the desired site, it is also critical to ensure that local irradiation is sufficient to drive the photoswitch and to suppress the symptoms of the disease, such as epileptiform activity (hippocampal paroxysmal discharges or HPDs) in a mouse TLE model. We first ascertained that Δ^9^−THC reduced the number of the HPDs in the chronic phase of the intrahippocampal kainic acid mouse model for refractory seizures. Next, we carried out exploratory experiments towards the first *in vivo* application of *azo*-THC-3, in which we directly infused the compound into the hippocampus, the site of seizure onset in the epileptic mice, combined with local irradiation (365 nm, 5 mW, 50 ms pulses at 10 Hz for 20 min).

In the positive control experiment, we found that intrahippocampal Δ^9^−THC (0.5 µg/0.5 µl) reduced the number of HPDs compared to vehicle (mixed effect analysis, treatment factor, F (1, 12) = 5.367; p = 0.0390) (Figure 3C). The reduction in hourly cumulative HPD time did however not reach statistical significance (mixed effect analysis, treatment factor, F (1, 3) =; 2.798 p = 0.1930) (Figure 3D). Because vehicle infusion caused a trend towards fewer HPD number and less HPD time, we refined the infusion protocol for subsequent experiments. This involved extending the handling period of the mice by at least two days and adjusting the setup to allow for drug delivery with minimal movement restriction to minimize stress effects on HPDs.

We next asked whether *in situ* photoactivation of *azo*-THC-3 (0.96 µg/0.5 µl) could suppress the spontaneous recurrent seizure discharges in chronically epileptic mice. *Azo*-THC-3 significantly reduced HPDs number, but only when irradiation at 365 nm had occurred (mixed effect analysis, treatment factor, F (1.673, 26.77) = 5.179; p = 0.0166) (Figure 3 B, E). A similar trend was observed for the hourly cumulative HPD time, but this did not reach statistical significance (mixed effect analysis, treatment factor, F (1.593, 11.15) = 4.166; p = 0.0516). (Figure 3 B, F).

### 3.4. Systemic injection and subsequent hippocampal *in situ* photoactivation of *azo*-THC-3 reduces the incidence of HPDs in the chronic intrahippocampal kainic acid mouse model

In a final experiment, we determined whether *in situ* photoactivating *azo*-THC-3 following its systemic administration could reduce HPDs in epileptic mice. HPDs were recorded during a baseline period of 2 h. Afterwards, Δ^9^−THC, *azo*-THC-3, or vehicle were injected which could then be followed by *in situ* irradiation (365 nm, 5 mW, 50 ms pulses at 10 Hz for 20 min) depending on the treatment group (Figure 4A). Δ^9^−THC was again used as a positive control. A systemic administration of Δ^9^−THC (10 mg/kg) nearly abolished HPDs. HPD number (mixed effect analysis, treatment factor, F (1.000, 6.000) = 8.626; p = 0.0260) and cumulative time (mixed effect analysis, treatment factor, F (1.000, 6.000) = 13.34; p = 0.0107) were significantly lower compared to vehicle (Figure 4 C, D). Notably, post-hoc testing revealed a significant reduction in the number of HPDs during the initial hour after injection (mean difference of - 56.57%, 95% CI: (−106.2, 6.905), p = 0.0304) and we observed significant differences in the hourly cumulative HPD time during both the first hour (mean difference of −58.60%, 95% CI: (−110.5, −6.698), p = 0.0315) and the second hour (−67.38%, 95% CI: (−125.1, −9.628), p = 0.0277) following injection. Finally, we evaluated the anticonvulsive potential of *azo*-THC-3 after systemic administration and in situ photoactivation. Mice were subjected to a total of 4 treatments (*azo*-THC-3 *in situ* irradiated with 365 nm light, *azo*-THC-3 pre-irradiated with 365 nm light, *azo*-THC-3 not irradiated, vehicle *in situ* irradiated with 365 nm light) again in a cross-over design with *ad random* order of the treatments. Here, pre-irradiated *azo*-THC-3 served as an additional control to evaluate the blood-brain barrier passage of the azobenzene containing molecule and as a point of reference for the efficacy of *in situ* irradiation. Administration of 23.9 mg/kg of *azo*-THC-3 pre-irradiated with 365 nm light exhibited (equivalent dose to Δ^9^−THC (10 mg/kg), compensating for the difference in molar mass and the incomplete conversion of *trans-azo*-THC tot *cis-azo-*THC) rapid anticonvulsant effects, manifesting within 5 to 20 min after injection (See supporting information). Based on this observation, we opted to initiate 20 min UV irradiation just 5 min following injection to photoactivate *azo*-THC-3 in *situ* at the level of the hippocampus (Figure 4 B, E, F). Significant differences were seen in the effects of the distinct treatments on HPD occurrence (mixed effect analysis, treatment factor, F (1.684, 16.84) = 10,51, p = 0.0016) and hourly cumulative time (mixed effect analysis, treatment factor, F (2.013,20.13) = 12.51, p = 0.0003). Importantly, when *azo*-THC-3 was injected while protected from light, without further *in situ* irradiation, it had no impact on HPD occurrence nor on hourly cumulative HPD time. Post-hoc analysis revealed that *in situ* irradiated *azo*-THC-3 significantly reduced HPD occurrence during the first hour after injection compared to the vehicle group (−66.96, 95% CI (−101.6, −32.31) adjusted p=0.0034) and compared to non-irradiated *azo*-THC-3 (−53.59, 95% CI (−95.44, −11.73) adjusted p=0.0196). Likewise, the hourly cumulative HPD time was significantly reduced during the first hour following injection ( - 57.65% (95% CI (−98.36, −16.94), adjusted p=0.0130)) and (−48.9% (95% CI (−88.00, −9.801), adjusted p=0.0216)) compared to the vehicle group and *azo*-THC-3 (not irradiated) group, respectively. The effect became less pronounced over the following time points. To account for any potential light-related effects on HPD occurrence, vehicle administration was also paired with *in situ* UV irradiation, but no discernible effect was noted.

### 3.5. Systemic injection and subsequent hippocampal *in situ* photoactivation of *azo*-THC-3 does not affect motor activity

A final key experiment should show whether our photoswitch approach for epilepsy with systemic administration of the molecule and *in situ* irradiation can also lead to fewer on-target side effects such as motor impairment. *Azo*-THC-3, pre-exposed to UV irradiation for 20 min before its systemic administration to the mice, led to overt changes in behaviour characterized by immobility over the entire 20-min time period (Figure 5A, ANOVA, treatment factor F (3, 18) = 9,521, p = 0.0005)). Crucially, immobility was not significantly different from the vehicle group when *azo*-THC-3 was non-irradiated or *in situ* photoactivated at the level of the hippocampus following its intraperitoneal delivery (Figure 5B, ANOVA, F (3, 18) = 9,521, p = 0.0005).

**Figure 5.**
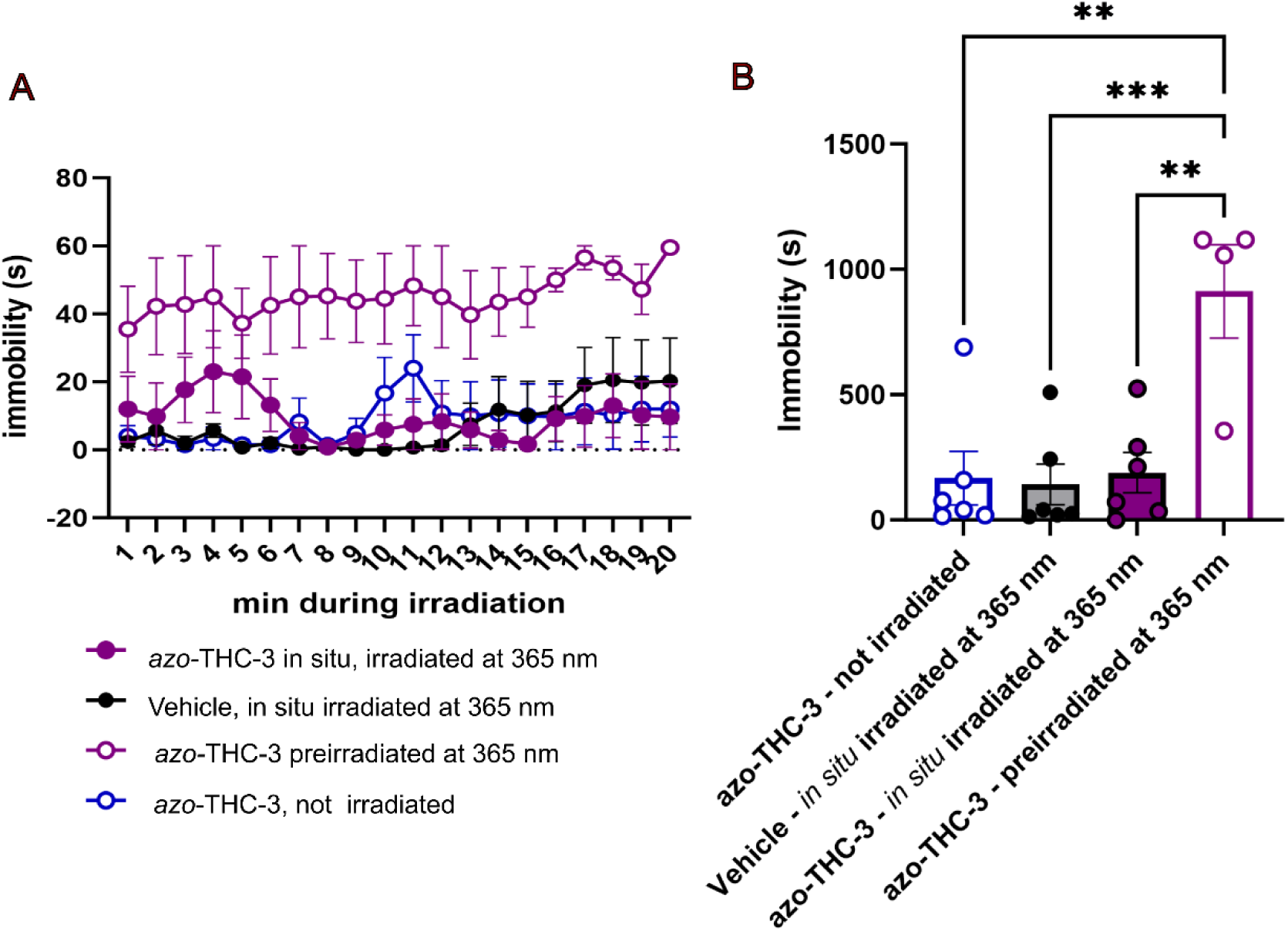
*Azo*-THC-3 *in situ* irradiated does not affect motor activity in a chronic TLE mouse model following systemic injection. (A) Time course of immobility measured in 1 min bins and (B) cumulative immobility during the 20 min irradiation period. Vehicle *in situ* irradiated at 365 nm group, n = 6; *azo*-THC-3 not irradiated group, n=6; *azo*-THC-3 pre-irradiated at 365 nm group, n= 4; *azo*-THC-3 *in situ* irradiated at 365 nm, n= 6. Data are presented as mean ± SEM. Overall treatment differences between experimental groups are indicated as ** p < 0.01 and *** p < 0.001.

To conclude, a photoswitchable Δ^9^−THC derivative, *azo*-THC-3, effectively decreased the burden of difficult-to-treat HPDs in chronic epileptic mice when *in situ* activated after systemic administration, without significantly impacting their locomotion.

## 4. Discussion

Temporal lobe epilepsy (TLE), a neurological disorder with a localised onset of seizures, is known as the most common cause of drug refractory epilepsy in adults^25^. First-line treatment of TLE typically entails pharmacological interventions using antiseizure medications that act upon voltage-gated ion channels, facilitate inhibitory neurotransmission or intervene with neuronal burst firing (e.g. via synaptic vesicle SV2A modulation). Nevertheless, about one-third of TLE patients keeps experiencing persistent seizures, due to low response to the currently available drugs or low tolerance for side effects^26–28^. Surgical resection of the seizure focus is an interesting, alternative option, but even in these patients, seizure recurrence after a period of seizure freedom is well described^29^. TLE with drug-refractory seizures thus remains a condition with a high, unmet, medical need for precise interventions that target the affected brain region to suppress seizures without leading to adverse effects in other parts of the body. Due to its focal onset, often the hippocampus, TLE may be well suited for photopharmacological intervention. Herein we present a novel strategy based on systemic administration of a photoswitchable Δ^9^−THC derivative, *azo*-THC-3, and its *in situ* photoactivation at the site of seizure onset in a mouse model of TLE with spontaneous difficult-to-treat seizures.

Δ^9^−THC is an intriguing molecule that is widely abused for its recreational properties but also gained considerable attention for possible medical applications. Cannabis-based treatments for epilepsy have a long history. The first indications date back to ancient times and case reports from the 19^th^ century support anticonvulsant effects of cannabis^30^. More recently, these anticonvulsant effects of cannabis-based medicinal products, notably cannabidiol, but also Δ^9^−THC, have been confirmed, especially for the treatment of refractory catastrophic epilepsies in children^31–34^. Large-scale clinical trials with Δ^9^−THC are lacking because of its psychoactive properties and the focus has shifted more to cannabidiol for epilepsy treatments^35^. However, Δ^9^−THC has been the subject of extensive testing in animal models for seizures and epilepsy^30^. Δ^9^−THC exhibited clear anticonvulsant effects in the acute maximal electroshock model for generalised seizures^36,37^ while results in the acute pentylenetetrazole seizure model were more variable (as reviewed in detail in Devinsky et al, 2024^30^). In epilepsy models, however, the effects of Δ^9^−THC seem more unambiguous but not yet test in the kainate mouse model. Δ^9^−THC possessed anticonvulsant effects against photogenic and amygdala-kindled seizures in baboons^38^, as well as in the rat amygdala kindling model^15^. Moreover, Δ^9^−THC abolished spontaneous recurrent generalised convulsions in the post-status epilepticus pilocarpine rat model^39^. However, in this paper the hippocampal refractory HPDs were not measured. We here confirm the anticonvulsive properties of Δ^9^−THC and its photoswitchable derivative *azo*-THC-3 in the mouse intrahippocampal kainic acid model for difficult-to-treat spontaneous epileptic discharges.

Finding an ideal photoswitchable molecule with translational value is a challenge. An important prerequisite is that the compound is little or not at all active at the time of its systemic administration, for applications in the field of neuroscience it is then important that the compound can cross the blood-brain barrier to a sufficient extent, and finally, after *in situ* irradiation at the desired site of action, the molecule should preferably be able to remain in its activated state for several min to generate appreciable pharmacodynamic effects. To the best of our knowledge, *azo*-THC-3 is a unique photoswitchable compound that meets these criteria. In contrast to the majority of GPCR-targeting photoswitches that have been published up to date^5,6^, *azo*-THC-3 is *cis*-active and bistable in aqueous environments. Upon exposure to UV light (365 nm), *azo*-THC-3 was previously shown to rapidly switch from the *trans* to *cis* configuration^17^. Importantly, *trans*-*azo*-THC-3 was reported to have a low potency (EC_50_ could not be determined accurately) and efficacy (E_max_ = 17%) for the CB1R. In contrast, *cis*-*azo*-THC-3 showed a high potency (EC_50_ = 17 nM) and sufficient efficacy (E_max_ = 52%) for the CB1R that were comparable to those of Δ^9^−THC^17^. We here confirm that the molecule switches from 13.9% ± 1.3% *cis*-*azo*-THC-3 in the absence of light to 71.7% ± 2.5% *cis*-*azo*-THC-3 after UV irradiation at 365 nm using previously reported pulsed irradiation conditions^22^. The obtained results align with previous reports of the azobenzene photoswitch properties^17^. The azobenzene motif is known for not supporting a full *trans*-to-*cis* conversion, with the 80% *cis* equilibrium representing the upper threshold for most compounds^5,40^. The *cis*-*azo*-THC-3 is relatively stable, with a half-life of 287 min for the spontaneous relaxation (at 37°C in the absence of light) to *trans*-*azo*-THC-3, again in line with previous observations^17^. The 10% *cis* equilibrium appears to correspond to the lower threshold under the used experimental conditions. Importantly, forced relaxation using blue light (450 nm), promoted a state in which 40% of the *cis*-*azo*-THC-3 remained present. This implies that the margin for testing rapid repeated switches between the active *cis*-*azo*-THC-3 and inactive *trans*-*azo*-THC-3 using UV and blue light irradiation parameters, respectively, that are tolerated by biological tissues is small, arguably too small for *in vivo* applications. Nevertheless, the extended relaxation half-life of *cis*-*azo*-THC-3 offered a significant advantage for our *in vivo* photopharmacological experiments in a mouse TLE model since it limited the need for extended illumination and frequent drug dosing.

In a first step, we demonstrated the protective action of *azo*-THC-3 in an acute *ex vivo* model for hippocampal hyperexcitability. We used Δ^9^−THC as a positive control, showing that it prevented the kainic acid-induced increase in EPSC frequency in murine hippocampal slices. This is in line with the notion that CB1R KO mice are more vulnerable to kainic acid-induced hyperexcitability^10^. Importantly, we found a similar protective effect against hyperexcitability for pre-irradiated *azo*-THC-3 but not for the non-irradiated *azo*-THC-3. As such, we established a proof-of-principle that when applied in the same concentration, photoswitching led to discernible biological effects of the *cis*- and the *trans*-*azo*-THC-3 isomers in a model for hippocampal hyperexcitability.

Next, we evaluated *in vivo* photoswitching in the hippocampus of chronically epileptic mice. We first infused the compounds directly into the hippocampus to address two potential issues of the photoswitching strategy for TLE. The strategy bypasses the blood-brain barrier and limits its therapeutic activity is mediated through the brain region that is targeted with irradiation. We demonstrated that the hippocampal seizure focus is indeed a viable target given that infusion of Δ^9^−THC had a powerful and long-lasting suppressive effect on the spontaneous recurrent HPDs in the post-status epilepticus kainic acid mouse model. These anticonvulsant effects of Δ^9^−THC on HPDs were not previously demonstrated. Moreover, we found that *azo*-THC-3 reduced the number of HPDs only when the infusion occurred simultaneously with *in situ* irradiation of 365 nm, while *azo*-THC-3 without irradiation or irradiation without *azo*-THC-3 had no significant effects on HPDs. This shows that *azo*-THC-3 can be switched *in vivo*, and that it can suppress refractory HPDs, on the condition that the hippocampal concentration of *cis*-*azo*-THC-3 is sufficiently high.

In the final experiment, we included also systemic injection of pre-irradiated *azo*-THC-3 as a positive control, which yielded a rapid and robust reduction of HPDs in the hippocampal formation. When non-irradiated *azo*-THC-3 was injected systemically and *in situ* irradiated we also found a significant reduction in HPD burden. As expected, *in situ* irradiation was slightly less efficient than pre-irradiation, corresponding to the notion that *in situ* irradiation can only convert the *azo*-THC-3 that eventually reaches the hippocampus and is located near the optical fibre. Nevertheless, the local conversion to *cis*-*azo*-THC-3 was sufficient to elicit a robust anti-seizure effect. Importantly, *azo*-THC-3 without irradiation or irradiation without *azo*-THC-3 had no significant effect on HPDs in these experiments.

Finally, we clearly demonstrate that systemic administration of pre-irradiated *azo*-THC-3 diminished locomotor activity in mice. It is well established that the equivalent dose of Δ^9^-THC causes hypolocomotion through activation of CB1Rs on principal neurons^41–43^. Additionally, exposure to pre-irradiated azo-THC-3 results in comparable motor impairments. Crucially, *in situ* hippocampal irradiation of *azo*-THC-3 had no significant effect on locomotor activity in mice, effectively demonstrating that the photoswitch strategy avoided a well-known on-target side effect of Δ^9^−THC

The used photoswitch strategy contrasts with photocaging, another primary strategy within the field of photopharmacology, where the active compound is attached to a photoprotective group that quenches the therapeutic effect. Upon irradiation, the photoprotective group is cleaved from the molecule and the initial properties of the drug are restored^44^. For example, ruthenium-bipyridine-triphenylphosphine γ−aminobutyric acid (RuBi-GABA), is a photocaged derivative of GABA that has shown to reduce ictal-like events in rat brains slices and terminated 4-aminopyridine-induced seizures *in vivo* after local application to the cortex^45^. While these results are very exciting, RuBi-GABA was reported to induce toxic effects at high doses^46^. In addition, the potential effects of the photocage itself, should be considered. Another example is the photocaged N^6^-cyclopentyladenosine (cCPA) that was used to achieve release of the adenosine A1 receptor agonist CPA using light of 405 nm thereby attenuating stimulus-evoked local field potentials in hippocampal brain slices^47^.

Regardless of the photopharmacological approach, it is imperative to account not only for properties of the drug itself, but also for the energy of the wavelength required to photoactivate the molecules. As such the effect of irradiation is a point of possible concern. The use of UV light in biological systems indeed presents inherent challenges to biocompatibility. Despite the wide adaptation of optogenetics in neuroscience for the last decade, the knowledge on the effect of UV(A) light in the brain specifically, remains limited. The total energy output in our 20 min irradiation protocol was 0.83 mWh. Previously, an irradiation dose of 2 mWh induced cortical oedema and triggered scar formation over 5 days post-exposure^48,49^. The extent to which irradiation parameters such as pulse duration, total irradiation duration, pulse frequency, and repeated exposure of brain structures affect tissue damage is unclear. In our present study we did not observe gross acute toxic effects of our irradiation protocol. Current evolutions in the photopharmacology field, such as red-shifted azobenzenes^50,51^, might be promising to reduce any potential tissue damage and to improve the depth of light penetration into the target tissue. Future advancements in red-shifted azobenzene applications and the development of wireless optic devices will facilitate the translation of our current photoswitchable azo-THC-3 strategy to clinical settings. In the proof-of-concept approach presented here, we still rely on UVA light delivered via an implanted optical fibre at the seizure focus, tethered to an external driver and light source.

In conclusion, this is the first report showing the successful application of a photoswitchable compound with systemic administration and local photoactivation in a brain region of interest. We thus provide an innovative proof-of-concept study for the application of photopharmacology, using a photoswitchable cannabinoid, to treat refractory focal seizures in TLE with unprecedented precision. Furthermore, we demonstrate that local activation of photoswitchable compounds may mitigate side effects associated with effective anti-seizure drugs.

## Acknowledgements

This research was funded by Research Foundation Flanders (FWO Vlaanderen), grant number 1136322N (PhD fellowship fundamental research S. Bournons), and by the senior research project G042219N (I. Smolders, D. De Bundel, Vrije Universiteit Brussel, and R. Raedt, University of Ghent). This research was also supported by the Vrije Universiteit Brussel (Strategic research programme SRP85). We would also like to express our sincere thanks to Surajit Sahu for his experimental contribution to this manuscript and would appreciate his updated contact details.

## Author information

S.B.: investigation, methodology, data collection, formal analysis, manuscript writing and editing

M.K.: investigation, methodology, project administration, manuscript review and editing.

B.K.: investigation, methodology, manuscript review and editing.

R.C.S.: conceptualization, investigation, methodology, manuscript review and editing.

E.H.: methodology, data collection, formal analysis, manuscript review and editing.

R.G.: methodology, manuscript review and editing.

P.P.: investigation, methodology, manuscript review and editing.

T.C.M.: methodology, manuscript review and editing.

M.V.W.: methodology, manuscript review and editing.

M.A.S.: methodology, manuscript review and editing.

G.D.S.: methodology, data collection and formal analysis

C.D.R: methodology, data collection and formal analysis

L.N.: methodology, manuscript review and editing

R.R.: conceptualization, funding acquisition, review and editing

E.M.C.: conceptualization, funding acquisition, supervision, manuscript review and editing.

D.D.B.: conceptualization, funding acquisition, supervision, manuscript writing, review and editing

I.S.: conceptualization, funding acquisition, supervision, manuscript writing, review and editing

## Data availability

Data available at: 10.5281/zenodo.13353702

## Conflict of interest

No conflict of interest to declare.

## Notes

### Competing Interest Statement

The authors have declared no competing interest.

## References

1 Thijs, R. D., Surges, R., O’Brien, T. J. & Sander, J. W. Epilepsy in adults. Lancet 393, 689–701 (2019). 10.1016/S0140-6736(18)32596-0

2 Cao, Z. et al. Progress in TLE treatment from 2003 to 2023: scientific measurement and visual analysis based on CiteSpace. Front Neurol 14, 1223457 (2023). 10.3389/fneur.2023.1223457

3 Karunakaran, S. et al. The interictal mesial temporal lobe epilepsy network. Epilepsia 59, 244–258 (2018). 10.1111/epi.13959

4 Vinti, V. et al. Temporal Lobe Epilepsy and Psychiatric Comorbidity. Front Neurol 12, 775781 (2021). 10.3389/fneur.2021.775781

5 Hull, K., Morstein, J. & Trauner, D. In Vivo Photopharmacology. Chem Rev 118, 10710–10747 (2018). 10.1021/acs.chemrev.8b00037

6 Pereira, V. & Goudet, C. Emerging Trends in Pain Modulation by Metabotropic Glutamate Receptors. Front Mol Neurosci 11, 464 (2018). 10.3389/fnmol.2018.00464

7 Velema, W. A., Szymanski, W. & Feringa, B. L. Photopharmacology: Beyond Proof of Principle. Journal of the American Chemical Society 136, 2178–2191 (2014). 10.1021/ja413063e

8 Frank, J. A. et al. In Vivo Photopharmacology Enabled by Multifunctional Fibers. ACS Chemical Neuroscience 11, 3802–3813 (2020). 10.1021/acschemneuro.0c00577

9 Ricart-Ortega, M., Font, J. & Llebaria, A. GPCR photopharmacology. Mol Cell Endocrinol 488, 36–51 (2019). 10.1016/j.mce.2019.03.003

10 Marsicano, G. et al. CB1 Cannabinoid Receptors and On-Demand Defense Against Excitotoxicity. Science 302, 84–88 (2003). doi:10.1126/science.1088208

11 Sugaya, Y. & Kano, M. Endocannabinoid-Mediated Control of Neural Circuit Excitability and Epileptic Seizures. Front Neural Circuits 15, 781113 (2021). 10.3389/fncir.2021.781113

12 Rosenberg, E. C., Tsien, R. W., Whalley, B. J. & Devinsky, O. Cannabinoids and Epilepsy. Neurotherapeutics 12, 747–768 (2015). 10.1007/s13311-015-0375-5

13 Cristino, L., Bisogno, T. & Di Marzo, V. Cannabinoids and the expanded endocannabinoid system in neurological disorders. Nat Rev Neurol 16, 9–29 (2020). 10.1038/s41582-019-0284-z

14 Rosenberg, E. C., Patra, P. H. & Whalley, B. J. Therapeutic effects of cannabinoids in animal models of seizures, epilepsy, epileptogenesis, and epilepsy-related neuroprotection. Epilepsy Behav 70, 319–327 (2017). 10.1016/j.yebeh.2016.11.006

15 Fallah, M. S., Dlugosz, L., Scott, B. W., Thompson, M. D. & Burnham, W. M. Antiseizure effects of the cannabinoids in the amygdala-kindling model. Epilepsia 62, 2274–2282 (2021). 10.1111/epi.16973

16 Stella, N. THC and CBD: Similarities and differences between siblings. Neuron 111, 302–327 (2023). 10.1016/j.neuron.2022.12.022

17 Westphal, M. V. et al. Synthesis of Photoswitchable Delta(9)-Tetrahydrocannabinol Derivatives Enables Optical Control of Cannabinoid Receptor 1 Signaling. J Am Chem Soc 139, 18206–18212 (2017). 10.1021/jacs.7b06456

18 Twele, F., Töllner, K., Brandt, C. & Löscher, W. Significant effects of sex, strain, and anesthesia in the intrahippocampal kainate mouse model of mesial temporal lobe epilepsy. Epilepsy & Behavior 55, 47–56 (2016). 10.1016/j.yebeh.2015.11.027

19 Laaris, N., Good, C. H. & Lupica, C. R. Delta9-tetrahydrocannabinol is a full agonist at CB1 receptors on GABA neuron axon terminals in the hippocampus. Neuropharmacology 59, 121–127 (2010). 10.1016/j.neuropharm.2010.04.013

20 Buckinx, A. et al. Translational potential of the ghrelin receptor agonist macimorelin for seizure suppression in pharmacoresistant epilepsy. Eur J Neurol 28, 3100–3112 (2021). 10.1111/ene.14992

21 Custers, M. L. et al. Neurofilament light chain: A possible fluid biomarker in the intrahippocampal kainic acid mouse model for chronic epilepsy? Epilepsia 64, 2200–2211 (2023). 10.1111/epi.17669

22 Zussy, C. et al. Dynamic modulation of inflammatory pain-related affective and sensory symptoms by optical control of amygdala metabotropic glutamate receptor 4. Mol Psychiatry 23, 509–520 (2018). 10.1038/mp.2016.223

23 Welzel, L., Schidlitzki, A., Twele, F., Anjum, M. & Loscher, W. A face-to-face comparison of the intra-amygdala and intrahippocampal kainate mouse models of mesial temporal lobe epilepsy and their utility for testing novel therapies. Epilepsia 61, 157–170 (2020). 10.1111/epi.16406

24 Widmann, M. et al. Characterization of the intrahippocampal kainic acid model in female mice with a special focus on seizure suppression by antiseizure medications. Exp Neurol 376, 114749 (2024). 10.1016/j.expneurol.2024.114749

25 Levesque, M. & Avoli, M. The kainic acid model of temporal lobe epilepsy. Neurosci Biobehav Rev 37, 2887–2899 (2013). 10.1016/j.neubiorev.2013.10.011

26 Devinsky, O. et al. Epilepsy. Nat Rev Dis Primers 4, 18024 (2018). 10.1038/nrdp.2018.24

27 Kwan, P. & Brodie, M. J. Definition of refractory epilepsy: defining the indefinable? Lancet Neurol 9, 27–29 (2010). 10.1016/S1474-4422(09)70304-7

28 Medel-Matus, J. S., Orozco-Suarez, S. & Escalante, R. G. Factors not considered in the study of drug-resistant epilepsy: Psychiatric comorbidities, age, and gender. Epilepsia Open 7 Suppl 1, S81–S93 (2022). 10.1002/epi4.12576

29 de Tisi, J. et al. The long-term outcome of adult epilepsy surgery, patterns of seizure remission, and relapse: a cohort study. Lancet 378, 1388–1395 (2011). 10.1016/S0140-6736(11)60890-8

30 Devinsky, O. A.-O., et al. Cannabinoid treatments in epilepsy and seizure disorders. (2024).

31 Gherzi, M. et al. Safety and pharmacokinetics of medical cannabis preparation in a monocentric series of young patients with drug resistant epilepsy. Complement Ther Med 51, 102402 (2020). 10.1016/j.ctim.2020.102402

32 Vickery, A. W. & Finch, P. M. Cannabis: are there any benefits? Intern Med J 50, 1326–1332 (2020). 10.1111/imj.15052

33 Nowicki, M. et al. Potential Benefit of Add-on Delta9-Tetrahydrocannabinol in Pediatric Drug-Resistant Epilepsy: A Case Series. Can J Neurol Sci 49, 595–597 (2022). 10.1017/cjn.2021.151

34 Erridge, S. et al. Clinical Outcome Data of Children Treated with Cannabis-Based Medicinal Products for Treatment Resistant Epilepsy-Analysis from the UK Medical Cannabis Registry. Neuropediatrics 54, 174–181 (2023). 10.1055/a-2002-2119

35 O’Connell, B. K., Gloss, D. & Devinsky, O. Cannabinoids in treatment-resistant epilepsy: A review. Epilepsy Behav 70, 341–348 (2017). 10.1016/j.yebeh.2016.11.012

36 Wallace, M. J., Wiley, J. L., Martin, B. R. & DeLorenzo, R. J. Assessment of the role of CB1 receptors in cannabinoid anticonvulsant effects. European Journal of Pharmacology 428, 51–57 (2001). 10.1016/S0014-2999(01)01243-2

37 Dlugosz, L., Zhou, H. Z., Scott, B. W. & Burnham, M. The effects of cannabidiol and Δ9-tetrahydrocannabinol, alone and in combination, in the maximal electroshock seizure model. Epilepsy Research 190, 107087 (2023). 10.1016/j.eplepsyres.2023.107087

38 Wada, J. A., Wake, A., Sato, M. & Corcoran, M. E. Antiepileptic and Prophylactic Effects of Tetrahydrocannabinols in Amygdaloid Kindled Cats*. Epilepsia 16, 503–510 (1975). 10.1111/j.1528-1157.1975.tb06080.x

39 Wallace, M. J., Blair, R. E., Falenski, K. W., Martin, B. R. & DeLorenzo, R. J. The endogenous cannabinoid system regulates seizure frequency and duration in a model of temporal lobe epilepsy. J Pharmacol Exp Ther 307, 129–137 (2003). 10.1124/jpet.103.051920

40 Bandara, H. M. & Burdette, S. C. Photoisomerization in different classes of azobenzene. Chem Soc Rev 41, 1809–1825 (2012). 10.1039/c1cs15179g

41 Monory, K. et al. Genetic dissection of behavioural and autonomic effects of Delta(9)-tetrahydrocannabinol in mice. PLoS Biol 5, e269 (2007). 10.1371/journal.pbio.0050269

42 Schreiber, S. et al. Functional effects of synthetic cannabinoids versus Delta(9) -THC in mice on body temperature, nociceptive threshold, anxiety, cognition, locomotor/exploratory parameters and depression. Addict Biol 24, 414–425 (2019). 10.1111/adb.12606

43 Han, Y. et al. Laminar Distribution of Cannabinoid Receptor 1 in the Prefrontal Cortex of Nonhuman Primates. Mol Neurobiol 61, 1–12 (2024). 10.1007/s12035-023-03828-4

44 Silva, J. M., Silva, E. & Reis, R. L. Light-triggered release of photocaged therapeutics - Where are we now? J Control Release 298, 154–176 (2019). 10.1016/j.jconrel.2019.02.006

45 Wang, D. et al. Photolysis of Caged-GABA Rapidly Terminates Seizures In Vivo: Concentration and Light Intensity Dependence. Front Neurol 8, 215 (2017). 10.3389/fneur.2017.00215

46 Yang, X., Rode, D. L., Peterka, D. S., Yuste, R. & Rothman, S. M. Optical control of focal epilepsy in vivo with caged gamma-aminobutyric acid. Ann Neurol 71, 68–75 (2012). 10.1002/ana.22596

47 Craey, E. et al. Ex Vivo Feedback Control of Neurotransmission Using a Photocaged Adenosine A1 Receptor Agonist. Int J Mol Sci 23 (2022). 10.3390/ijms23168887

48 Nakata, M., Nagasaka, K., Shimoda, M., Takashima, I. & Yamamoto, S. Focal brain lesions induced with ultraviolet irradiation. Sci Rep 8, 7968 (2018). 10.1038/s41598-018-26117-w

49 Nakata, M., Shimoda, M. & Yamamoto, S. UV-Induced Neuronal Degeneration in the Rat Cerebral Cortex. Cereb Cortex Commun 2, tgab006 (2021). 10.1093/texcom/tgab006

50 Dong, M., Babalhavaeji, A., Samanta, S., Beharry, A. A. & Woolley, G. A. Red-Shifting Azobenzene Photoswitches for in Vivo Use. Acc Chem Res 48, 2662–2670 (2015). 10.1021/acs.accounts.5b00270

51 Zhang, M. et al. TRP (transient receptor potential) ion channel family: structures, biological functions and therapeutic interventions for diseases. Signal Transduct Target Ther 8, 261 (2023). 10.1038/s41392-023-01464-x

